# Variability in carbapenemase activity of intrinsic OxaAb (OXA-51-like) beta-lactamase enzymes in *Acinetobacter baumannii*

**DOI:** 10.1101/2020.07.02.183822

**Authors:** Yuiko Takebayashi, Jacqueline Findlay, Kate J. Heesom, Philip J. Warburton, Matthew B. Avison, Benjamin A. Evans

**Author notes:** Correspondence: Yuiko Takebayashi, School of Cellular and Molecular Medicine, Biomedical Sciences Building, University Walk, Bristol, UK, BS8 1TD.

## Abstract

**Objectives:** This study aimed to measure the variability in carbapenem susceptibility conferred by different OxaAb variants, characterise the molecular evolution of *oxaAb* and elucidate the contribution of OxaAb and other possible carbapenem resistance factors in the clinical isolates using WGS and LC-MS/MS.

**Methods:** Disc susceptibility and MIC broth microdilution tests were performed on ten clinical *A. baumannii* isolates and interpreted according to CLSI guidelines. Imipenem and meropenem MICs were evaluated for all *oxaAb* variants cloned into susceptible *A. baumannii* CIP70.10 and increased *adeABC* efflux pump gene expression BM4547 backgrounds, with and without their natural promoters. Molecular evolution analysis of the *oxaAb* variants was performed using FastTree and SplitsTree4. Resistance determinants were studied in the clinical isolates using WGS and LC-MS/MS analysis.

**Results:** OxaAb(82), OxaAb(83), OxaAb(107), and OxaAb(110) carrying I129L and L167V substitutions increased carbapenem MICs when transferred into susceptible *A. baumannii* backgrounds without an upstream IS element. Carbapenem resistance was conferred with the addition of their natural upstream IS*Aba1* promoter. LC-MS/MS analysis on the original clinical isolates showed no differences in expression levels of proteins commonly associated with carbapenem resistance.

**Conclusions:** IS*Aba1*-driven overexpression of OxaAb variants with substitutions I129L and L167V confers carbapenem resistance with no need for additional resistance mechanisms.

## INTRODUCTION

Carbapenem-resistant *Acinetobacter baumannii* is a World Health Organisation (WHO) priority level one pathogen, commonly associated with nosocomial infections in intensive care units (ICUs).^1,2^ Once treatable with broad-spectrum cephalosporins such as ceftazidime and cefepime, heavy usage of these antibiotics has led to the reliance and subsequent resistance to last-resort carbapenem treatment. *A. baumannii* are notorious for their genetic plasticity, enabling them to acquire resistance genes from *Pseudomonas aeruginosa* and clinically relevant Enterobacteriaceae species such as *Escherichia coli* and *Klebsiella pneumoniae*. Clinical *A. baumannii* have been reported to carry multiple acquired β-lactamases from all four Class A-D molecular groups such as TEM, CARB, PER, GES, VEB, CTX-M, IMP, VIM, NDM and OXA to varying frequencies, in addition to the intrinsic AmpC (ADC) and OxaAb (OXA-51-like) enzymes.^3–6^ Upregulation of some of these β-lactamases by means of insertion sequences (IS) such as IS*Aba1* and IS*Aba125* have also driven this resistance phenomenon.^7–9^

The main mechanism for carbapenem resistance in *A. baumannii* is carbapenem-hydrolysing class D β-lactamases, most commonly Oxa23, Oxa40, OxaAb, Oxa58, Oxa143 and Oxa235 groups, frequently associated with IS elements.^10–15^ Characterisation of clinical isolates has also inferred the synergistic importance of the upregulation of multidrug efflux pumps (notably the RND transporters AdeABC and AdeIJK) and the loss of certain porins (CarO, Omp33-36, OmpA and OmpW).^16–22^ However, the extent in which these proteins and their production levels play a role in carbapenem resistance is not yet clear. In recent years there has been a concerted effort to fill these gaps in our understanding of the factors contributing to carbapenem resistance phenotypes in *A. baumannii* using WGS, whole transcriptome shotgun sequencing (RNA-Seq) and proteomic approaches. However, these studies have not always been consistent with one another - in some cases carbapenem-resistant strains were shown to overexpress efflux pumps and downregulate porins^17,20^, whereas in others, carbapenem resistance was associated with an increase in porin abundance.^18^ These inconsistencies demonstrate that our understanding of the interplay between resistance mechanisms in *A. baumannii* remains incomplete.

OxaAb enzymes are intrinsic and by far the largest group of OXAs in *A. baumannii*, with 320 variants identified as of 03/06/2020.^23^ When OxaAb variants are characterised in clinical carbapenem resistant isolates worldwide, the presence of IS*AbaI* upstream is frequently noted and this has led to the general acceptance that transcriptional upregulation of these enzymes by upstream IS insertion, providing a strong promoter, can confer carbapenem resistance in the absence of other β-lactamases. However, it is unclear whether only specific variants (e.g. OxaAb(138) and OxaAb(82)^24,25^) confer this phenotype or if overproduction of all OxaAb types can lead to carbapenem resistance. Studies from the last few years of the effect of specific amino acid substitutions in OxaAb, for example at Ile-129, Leu-167 and Trp-222, have demonstrated that this can alter the enzyme structure and significantly increase catalytic activity with respect to the carbapenems.^26–29^ However, the impact of such substitutions alone on the antibiotic susceptibility of bacteria is unclear. Recent papers have highlighted clinical isolates carrying IS*Aba1*/*oxaAb* genes that do not exhibit carbapenem resistance.^30,31^ Nigro and Hall also elude to differences in carbapenem MIC depending on the OxaAb variant and/or other intrinsic factors in different backgrounds.^31^ In order to address some of these stated unknowns concerning carbapenem resistance in *A. baumannii*, this study aimed to i) measure the variability in carbapenem MIC conferred by different OxaAb variants, ii) characterise the molecular evolution of *oxaAb* and iii) elucidate the contribution of OxaAb and other possible carbapenem resistance determinants in clinical isolates using WGS and LC-MS/MS proteomics.

## MATERIALS AND METHODS

### Bacterial strains and antimicrobial susceptibility testing

Ten clinical *A. baumannii* isolates were used in this study (**Table 1**). Imipenem and meropenem MICs were previously characterised by Evans *et al* (except isolates B1 and A403), as well as the identification of IS*AbaI* elements upstream of their respective *oxaAb* genes.^32^ Recombinants were made using the following strains: *E. coli* DH5α (Subcloning Efficiency DH5α Competent cells, Invitrogen, United Kingdom), *A. baumannii* CIP70.10 and BM4547^33^ (gifts from Laurent Poirel, University of Fribourg). Disc susceptibility and MIC broth microdilution tests were performed and interpreted according to CLSI guidelines.^34^

**Table 1.** Selected WGS data and carbapenem minimal inhibitory concentrations (MIC) of clinical *A. baumannii* isolates.

| *A. baumannii*<br>ID | Geographic<br>Location | OxaAb | Global<br>Clone (GC) | Other β-lactamases | IS*Aba1*<br>OXA | IS*Aba1*<br>ADC | IS*Aba125*<br>ADC | Tn3<br>TEM | MIC (mg/L) | |
| --- | --- | --- | --- | --- | --- | --- | --- | --- | --- | --- |
|  |  |  |  |  |  |  |  |  | IMP | MEM |
| CIP70.10 | France | (64) | - | ADC-50 | - | - | - | - | 0.125-0.25 | 0.125-0.5 |
| A37 | Singapore | (64) | - | ADC-174 | - | + | - | - | 0.5 | 0.5 |
| A60 | Argentina | (65) | - | ADC-5, TEM-1A, CARB-16 | - | + | - | + | 0.125 | 2 |
| A230 | United Kingdom | (66) | 1 | ADC-175, Oxa20 | - | - | + | - | 0.5 | 2 |
| A90 | United Kingdom | (69) | 2 | ADC-11, TEM-1D | - | - | - | + | 0.125 | 0.25 |
| B1 | Unknown | (51) | - | ADC-180, Oxa10 | - | - | - | - | 1 | 1 |
| A403 | Taiwan | (82) | 1 | ADC-177, TEM-1D | + | + | - | + | 32 | 32 |
| A371 | Czech Republic | (83) | 1 | ADC-30, TEM-1D | + | + | - | + | 16 | 32 |
| A443 | Slovenia | (107) | 2 | ADC-176, TEM-1D | + | + | - | + | 16 | 16 |
| A404 | Poland | (110) | 2 | ADC-178, TEM-1D | + | + | - | + | 8 | 16 |
| A135 | Belgium | (111) | 2 | ADC-179 | - | - | - | - | 0.25 | 0.25 |

Resistant IMP and MEM MIC values ( $\geq 8$ mg/L) shaded grey. IS*Aba1*, IS*Aba125* and Tn3 sequences were found upstream of *oxaAb*, *bla*<sub>ADC</sub> or *bla*<sub>TEM</sub> genes. IMP, imipenem; MEM, meropenem.

### Whole genome sequencing (WGS)

Genomes were sequenced by MicrobesNG on a HiSeq 2500 instrument (Illumina, San Diego, CA, USA) as previously described.^35^ Insertion Sequences (IS) were identified using ISFinder.^36^

### Proteome analysis via Orbitrap LC-MS/MS

Total cell extractions of the clinical isolates (in three biological replicates) were prepared and 1 μg of each sample was analysed using an Orbitrap Velos mass spectrometer (Thermo Fisher Scientific) and quantified using Proteome Discoverer software v1.4 (Thermo Fisher Scientific) as outlined previously.^35^ The raw data files were searched against the UniProt *A. baumannii* ACIBA database (67,615 protein entries) and an in-house mobile resistance determinant database.^37^ Abundance values of each protein were converted to ratios relative to the average abundance of 30S and 50S ribosomal proteins, for ease of comparison between isolates.

### Cloning genes, transformation and sequencing

The genes for the *oxaAb* variants encoding OxaAb(64), (65), (66), (69), (71), (82), (83), (107), (110) and (111) were PCR amplified from clinical *A. baumannii* isolates (with additional NcoI and XhoI sites introduced at the 5’ and 3’ ends respectively) using OXA-66-NcoI F, OXA-111-NcoI F or OXA-71-NcoI F and OXA-66-XhoI R primers (**Table 2**) and TA cloned into the vector pGEM-T Easy (Promega, United Kingdom). The inserts were confirmed by sequencing with the universal T7 Promoter primer. For transformation into *E. coli* DH5α, *A. baumannii* CIP70.10 and BM4547, the inserts were digested with NcoI and XhoI and ligated into pYMAb2 (a gift from Dr Te-Li Chen, National Defense Medical Center, Taiwan). For genes including their natural upstream promoter regions, inserts were PCR amplified using OXA-51-like_XbaI F or ISAba1_XbaI F and OXA-51-like_EcoRI R primers, digested with XbaI and EcoRI and ligated into pUBYT.^37^ All inserts were confirmed by sequencing using the pYMAb2 Check primers (**Table 2**).

**Table 2.**
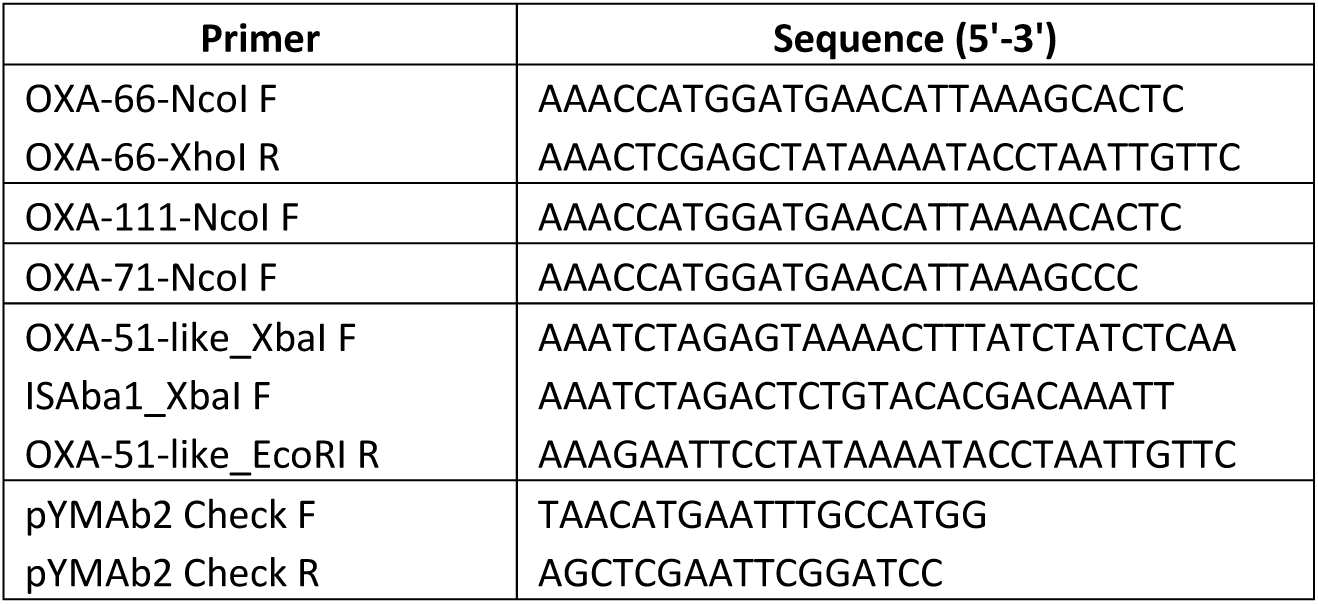
Primers used in this study.

All plasmids were used to transform *E. coli* DH5α and *A. baumannii* CIP70.10 and BM4547 strains by electroporation. Transformants were selected with ampicillin (100 mg/L) and ChromoMax IPTG/X-Gal (Fisher BioReagents, United Kingdom) for pGEM-T Easy recombinants or kanamycin (50 mg/L) for pYMAb2 and pUBYT recombinants.

### Predicting the molecular evolution of OxaAb variants

The nucleotide sequence of *oxaAb(66)* was used to query the NCBI nucleotide and genome databases using BLAST, implemented in Geneious (https://www.geneious.com), and all available *oxaAb* sequences were downloaded. The sequence for the gene of the naturally-occurring OXA from *Acinetobacter calcoaceticus* (*oxa213*) was included as an outgroup.^38^ Duplicate sequences were removed and remaining sequences aligned. A maximum likelihood phylogeny of the *oxaAb* genes was estimated using FastTree. Support for the resulting phylogeny was estimated using 100 bootstraps. The Phi test was used to detect recombination within the *oxaAb* alignment using SplitsTree4.^39^ A translation of the nucleotide alignment was used to identify all OXAs that were different from the consensus sequence at Ile-129 and Leu-167 that have previously been described as being important for substrate specificity and hydrolytic activity.^26,27,40^

## RESULTS AND DISCUSSION

### Characterisation of β-lactam susceptibility in selected clinical isolates

Ten clinical isolates^32^ (**Table 1**) were chosen for encoding various OxaAb enzymes that are representative of global clones (GC) 1 and 2 (OxaAb(69) and OxaAb(66) respectively). Some of these isolates (A371, A404, A403 and A443) were also chosen for encoding variants containing substitutions at sites considered important for substrate specificity (I129L in OxaAb(83) and OxaAb(110), and L167V in OxaAb(82) and OxaAb(107) respectively). Others (A60, A37 and A135) were chosen to represent polymorphisms at sites previously identified as having arisen more than once across the OxaAb phylogeny (Glu-36 in OxaAb(65), and Q194P in OxaAb(64) and OxaAb(111) respectively).^26,27,40^

β-lactam susceptibility results for the clinical isolates are shown in **Table 3**. All isolates were not susceptible to ceftriaxone and cefotaxime except for A135. The four isolates encoding variants with substitutions I129L and L167V (A404, A371, A443 and A403) were non-susceptible to all tested antibiotics, including the carbapenems. **Table 1** summarises the carbapenem MICs and the designation of IS elements upstream of β-lactamase genes based on the WGS data. Apart from the *oxaAb* variants, WGS did not detect any other β-lactamases known to confer carbapenem resistance.

**Table 3.** Disc susceptibility test results for selected β-lactams.

| ***A. baumannii* ID** | **CRO** | **CTX** | **CAZ** | **FEP** | **IPM** | **MEM** | **DOR** |
| --- | --- | --- | --- | --- | --- | --- | --- |
| CIP70.10 | I | I | S | S | S | S | S |
| A37 | R | R | R | S | S | S | S |
| A60 | R | R | I | R | S | S | S |
| A230 | R | R | R | I | S | S | S |
| A90 | I | R | S | S | S | S | S |
| B1 | I | I | S | S | S | S | S |
| A403 | R | R | R | R | R | R | R |
| A371 | R | R | R | R | R | R | R |
| A443 | R | R | R | R | R | R | R |
| A404 | R | R | R | I | R | R | R |
| A135 | S | I | S | S | S | S | S |

R, resistant; I, intermediate; S, susceptible; CRO, ceftriaxone; CTX, cefotaxime; CAZ, ceftazidime; FEP, cefepime; IPM, imipenem; MEM, meropenem; DOR, doripenem.

### OxaAb variants OxaAb(82), (83), (107), and (110) increase carbapenem MICs

To determine whether substitutions at Ile-129 and Leu-167 in OxaAb contribute to the observed carbapenem resistance in isolates A404, A371, A443 and A403, all *oxaAb* genes from the 10 clinical isolates were cloned in the absence of their native promoter, all downstream of the same Oxa24(72) promoter carried by pYMAb2. This was to exclude any confounding effects on differential gene expression of upstream IS elements seen in the clinical isolates.

In an *A. baumannii* CIP70.10 background (representing a susceptible host), OxaAb variants with a substitution at either Ile-129 or Leu-167 allowed for significantly increased carbapenem MICs over the other OxaAb variants (t-test, meropenem: *p* = 0.0196, imipenem: *p* = 0.0131) (**Table 4**). The same was true in the *A. baumannii* BM4547 background, which has increased *adeABC* efflux pump gene expression^33^ (t-test, meropenem: *p* = 0.0239, imipenem: *p* = 0.0391). Only OxaAb(82) conferred meropenem resistance (8 mg/L) in both backgrounds. The presence of OxaAb(107) and (83) increased meropenem MIC to an intermediate phenotype (4 mg/L) in both CIP70.10 and BM4547. Interestingly there was no overall difference in the MIC values between the CIP70.10 and BM4547 backgrounds (t-test, meropenem: *p* = 0.9209, imipenem: *p* = 0.6887). We therefore conclude that the Ile-129 or Leu-167 substitutions seen in OxaAb(82), (83), (107) and (110), increase carbapenem MIC but not to the level of resistance seen in the clinical isolates producing these variants. Furthermore, AdeABC overproduction is not important for carbapenem MICs in strains producing these OxaAb variants.

**Table 4.** Minimal inhibitory concentration (MIC in mg/L) of recombinant *A. baumannii* strains carrying various OxaAb enzymes ± their natural upstream promoter regions.

| Strain | pYMAb2 |  | pUBYT |  |
| --- | --- | --- | --- | --- |
|  | IPM | MEM | IPM | MEM |
| **CIP70.10 (No Vector)** | 0.125 | 0.125 | 0.125 | 0.125 |
| Empty Vector | 0.25 | 0.25 | 0.125 | 0.25 |
| OxaAb(64) | 0.125 | 0.5 | 0.5 | 4 |
| OxaAb(65) | 0.125 | 0.125 | 0.5 | 4 |
| OxaAb(66) | 0.25 | 0.25 | 0.25 | 4 |
| OxaAb(69) | 0.125 | 0.125 | 0.25 | 4 |
| OxaAb(51) | 0.06 | 0.25 | 0.25 | 4 |
| OxaAb(82)* | 2 | 8 | 16 | 64 |
| OxaAb(83)* | 0.5 | 2 | - | - |
| OxaAb(107)* | 2 | 4 | 16 | 64 |
| OxaAb(110)* | 0.5 | 2 | 8 | 64 |
| OxaAb(111) | 0.25 | 0.125-0.5 | 0.25 | 1 |
| **BM4547 (No Vector)** | 0.125 | 0.5 | 0.125 | 0.5 |
| Empty Vector | 0.125 | 0.5 | 0.125 | 0.25 |
| OxaAb(64) | 0.25 | 0.125-0.5 | 0.5 | 4 |
| OxaAb(65) | 0.125 | 0.5 | 0.5 | 4 |
| OxaAb(66) | 0.25 | 0.5 | 1 | 8 |
| OxaAb(69) | 0.125 | 0.5 | 0.5 | 2 |
| OxaAb(51) | 0.25 | 0.5 | 0.5 | 4 |
| OxaAb(82)* | 2 | 8 | 32 | > 64 |
| OxaAb(83)* | 0.5 | 4 | - | - |
| OxaAb(107)* | 1 | 2 | 64 | > 64 |
| OxaAb(110)* | 0.25 | 2 | 32 | > 64 |
| OxaAb(111) | 0.06 | 0.125 | 0.125 | 4 |

IPM, imipenem; MEM, meropenem. Intermediate (4 mg/L) and resistant (≥ 8 mg/L) MIC values shaded grey. “-” indicates strains were not tested. MIC values (n=6) that were variable are represented by ranges. “\*” highlight OxaAb variants with substitutions in Ile-129 or Leu-167.

### ISAbaI-driven expression of oxaAb only confers carbapenem resistance for certain oxaAb variants

The *oxaAb* variants were next cloned into pUBYT with their natural upstream promoter, to identify if the presence of upstream IS elements can enhance MICs and confer carbapenem resistance. Genes encoding three enzymes (OxaAb(82), (107), (110)) with changes at positions Ile-129 or Leu-167, had natural promotors provided by IS*Aba1*, while the remaining *oxaAb* genes had the native chromosomal promotor without the presence of an insertion sequence. Transformation of OxaAb(83) was not achieved despite multiple attempts.

When expressed in CIP70.10, the carbapenem MICs against transformants encoding *oxaAb* with an IS*Aba1* promotor increased to resistant levels seen in their parent clinical isolates (**Table 4**). These were significantly higher than the MICs obtained for the other *oxaAb* variants (t-test, meropenem – *p* = 9.99×10^−12^, imipenem – *p* = 1.45×10^−4^) where clinical resistance was not reached. While there was an overall increase in meropenem MICs for all transformants under the control of their native promoters compared to the pYMAb2 promoter (t-test, meropenem – p = 0.0465, imipenem – p = 0.0817), significantly higher MICs were observed for the three transformants encoding *oxaAb* with an IS*Aba1* promoter (t-test, meropenem – *p* = 0.0009, imipenem – *p* = 0.0319). This demonstrates that the addition of IS*AbaI* upstream of *oxaAb* variants encoding I129L or L167V substitutions confers carbapenem resistance in a recombinant background without any other resistance determinants. When the *oxaAb* variants were cloned into BM4547, the same pattern was also observed and there was no overall difference in the MIC values between the CIP70.10 and BM4547 backgrounds (t-test, meropenem: p = 0.9700, imipenem: p = 0.2391). Therefore, the increase in AdeABC efflux does not have a crucial role in conferring carbapenem resistance in the context of these OxaAb variants.

It is worth noting that the recombinants with upstream IS*Aba1* were very difficult to obtain, with extremely low transformation efficiency in both CIP70.10 and BM4547. Plasmid-mediated carriage of these variants with upstream IS*Aba1* may be deleterious to the host’s fitness and may possibly be the reason for certain variants not being observed to be plasmid-borne in nature.

### Predicting molecular evolution of oxaAb

Given that OxaAb variants with specific amino acid polymorphisms at Ile-129 and Leu-167 have been shown to confer carbapenem resistance in the presence of IS*Aba1*, it is reasonable to hypothesise that these polymorphisms may have been selected for in the *A. baumannii* population. To investigate the distribution of these two polymorphisms, a phylogenetic analysis of all available *oxaAb* genes was conducted. Comparison of the *oxaAb* phylogeny with the substitution patterns that result in amino acid changes at the two sites examined showed that substitutions at these sites are likely to have occurred on multiple occasions (**Figure 1, Table S1**). At position 129, there are 4 alternative codons coding for 4 amino acid changes, suggesting at least 4 independent mutations at this position. The phi test did not detect any significant evidence for recombination within the *oxaAb* genes. Therefore, assuming there is no recombination within these alleles, their distribution across the *oxaAb* phylogeny indicates that mutations at position 129 have occurred on 10 occasions, as seen by alleles carrying the same mutation being separated by alleles that do not share the mutation. Similarly, 3 amino acid changes at position 167 are coded for by 4 different codons, with a phylogenetic distribution suggesting independent mutations arising on 8 occasions. Overall, these data provide strong evidence for selection for changes to the consensus sequence at these sites. Given that no evidence for recombination within the *oxaAb* genes was detected, there are two possibilities that may explain the distribution of polymorphisms: 1) all of the variants have evolved independently and in some instances represent parallel evolution, and 2) there has been recombination of entire *oxaAb* genes between strains, most likely by natural transformation. The most conservative interpretation would be that each different codon only evolved once and any occurrences of the same codon are due to common evolutionary descent or recombination. While we did not detect evidence for recombination within the *oxaAb* genes here, the possibility of between-strain recombination could be examined by a large whole genome analysis, provided sufficient representation of the different *oxaAb* variants were included. At the other extreme, the most liberal interpretation of the data is that each different codon has evolved independently except where there is common evolutionary descent. The relative contributions of independent mutation and recombination to the evolutionary genetics of *oxaAb* remains to be determined; however, our experimental data does show that selection for changes in these two sites do increase carbapenem MIC.

**Figure 1.**
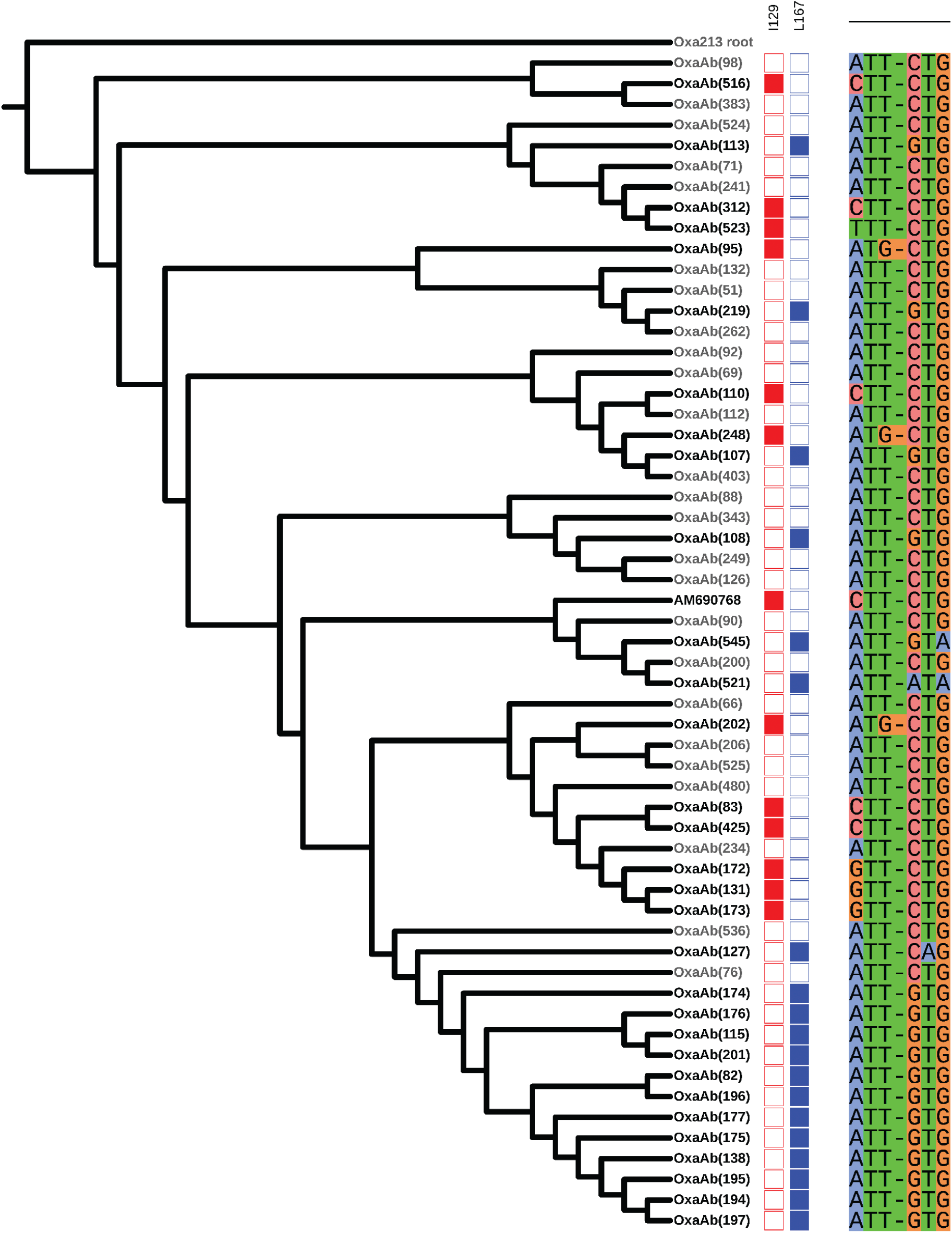
Cladogram of the nucleotide phylogeny of selected *oxaAb* sequences and their differences from consensus at amino acid positions 129 and 167. The phylogeny was drawn using FastTree with all available *oxaAb* sequences and rooted using *oxa213* from *Acinetobacter calcoaceticus* as an outgroup. The sequence labelled accession number AM690768 is an unnamed variant differing from OxaAb(90) by a single amino acid (at position 129). For clarity, the majority of branches containing genes for OxaAb enzymes that do not have a change from consensus at either position being examined have been hidden, with a minority retained to provide context (shown in italic font). The boxes in the centre represent the amino acid positions, with changes from consensus represented by a filled box. On the right are shown the sequences of amino acids and the corresponding codons. The figure was drawn using iTOL^57^.

### Comparing carbapenem resistance signatures in clinical isolates by LC-MS/MS and WGS

To determine if upregulation of OxaAb with substitutions in Ile-129 and Leu-167 is the only mechanism of carbapenem resistance in the four resistant clinical isolates (A403, A371, A443 and A404), all 10 clinical isolates were analysed for the presence of known carbapenem resistance proteins that may be differentially produced in the resistant isolates. Whole cell extracts were analysed in triplicate via LC-MS/MS and compared against each other and the reference strain CIP70.10. Over 2,000 proteins were identified and from these, abundance levels of porins, efflux pumps and β-lactamases commonly associated with carbapenem resistance are summarised in **Figure 2**.

**Figure 2.**
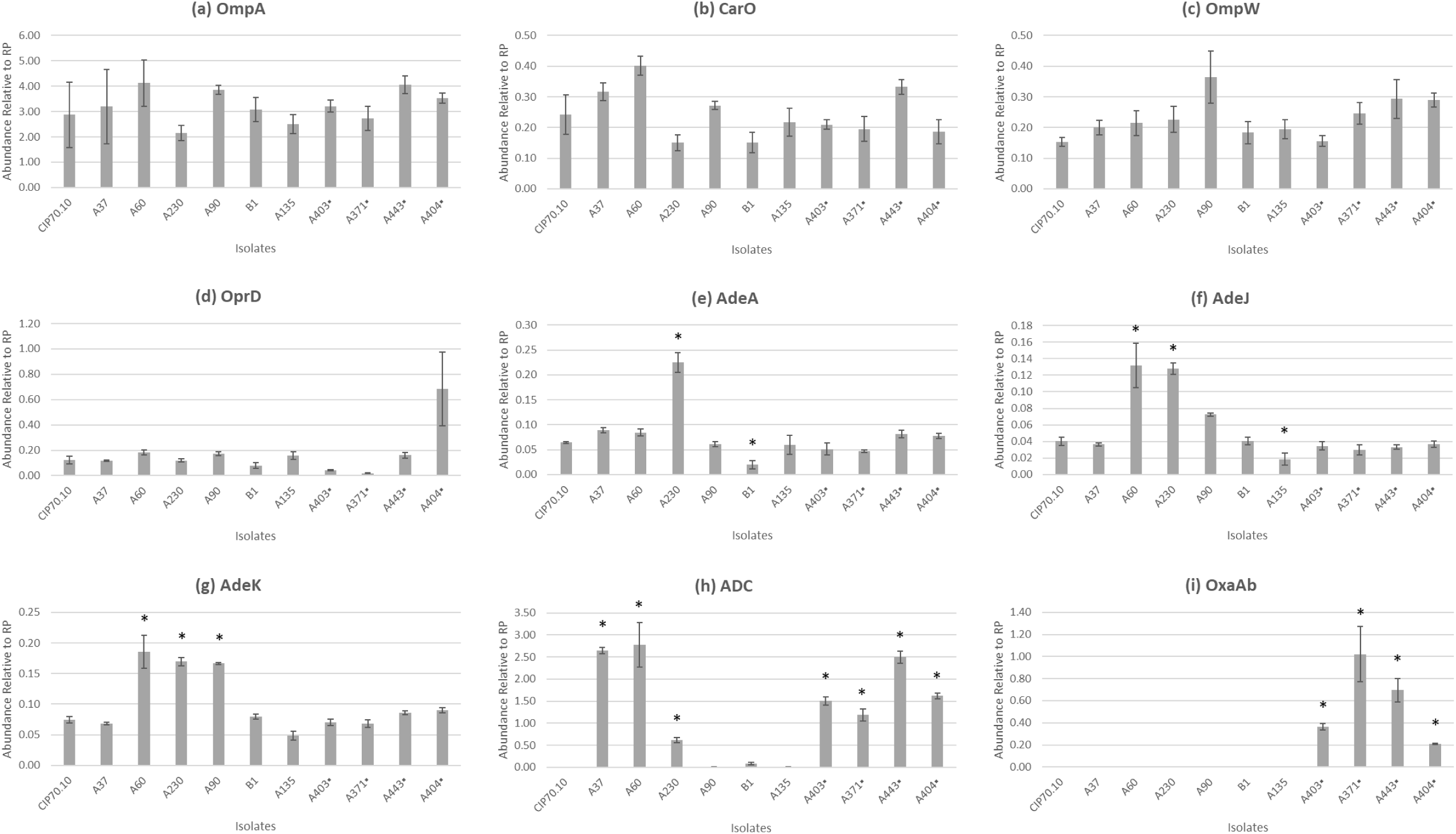
Comparison of various resistance determinants by average abundance ratios relative to ribosomal protein (RP). **(a)** OmpA, **(b)** CarO, **(c)** OmpW, **(d)** OprD, **(e)** AdeA, **(f)** AdeJ, **(g)** AdeK, **(h)** ADC and **(i)** OxaAb enzymes. Carbapenem resistant clinical isolates are highlighted with “▪”. The absolute abundance values of each protein of interest were divided by the average abundance values of 30S and 50S ribosomal proteins and averaged to yield ratios with SEM error bars (n=3). Asterisks represent a significant difference in abundance relative to CIP70.10, based on ≥ 2-fold difference and t-test (p<0.05).

#### (i) Porins

In *A. baumannii*, the major outer membrane protein associated with antimicrobial resistance is a nonspecific slow porin OmpA.^41^ It is generally accepted that OmpA is involved in the slow diffusion of certain β-lactams across the membrane.^19,41,42^ There were no changes in abundance levels of OmpA in the clinical isolates (compared to CIP70.10) (**Figure 2a**).

In terms of porins associated with carbapenem susceptibility, abundance levels of CarO, OmpW and OprD were compared. The disruption of CarO expression in MDR clinical *A. baumannii* isolates by insertion sequences (such as IS*Aba1*, IS*Aba10*, IS*Aba125* and IS*Aba825*) has been associated with reduced susceptibility to imipenem^16,43^, although a reconstituted liposome CarO system has been demonstrated not to transport imipenem.^44^ CarO can be grouped into two major isoforms CarOa and CarOb, with higher specificity for imipenem in the latter.^17^ WGS identified all isolates to have intact *carO* genes and no changes to the upstream promoter sequences. A60 and B1 carry CarOa and all other isolates carry CarOb, except A37, A443 and A135 which did not categorise in either groups. Abundance levels of CarO were similar across all isolates except A60, suggesting that there is no critical association between production levels or specific isoforms and carbapenem MIC in these clinical isolates (**Figure 2b**).

Loss of OmpW has been implicated with carbapenem resistance, although proteomics studies have also observed increased levels of this porin in MDR isolates.^21^ Another study observed that deletion of *ompW* in carbapenem-susceptible *A. baumannii* ATCC 17978 did not affect imipenem MIC.^45^ No differences in abundance levels were observed (except A90), suggesting that OmpW does not play a role in carbapenem resistance in these isolates (**Figure 2c**). Decreased *oprD* expression has been associated with carbapenem resistance in clinical isolates,^46–48^ although subsequent knock-out experiments demonstrated no increase in imipenem and meropenem susceptibilities.^49,50^ More recent liposome model studies have shown that OprD does uptake both carbapenems.^44^ Here we observed no significant changes in OprD abundance levels compared to CIP70.10 (**Figure 2d**).

#### (ii) Efflux pumps

Overexpression of RND efflux pumps AdeABC and AdeIJK have been associated with aiding carbapenem resistance, although this was not the case in our BM4547 recombinants.^51–53^ AdeB was below the level of detection in CIP70.10 and A135, despite confirmation of the gene by WGS. AdeC was not detected in CIP70.10, A37, A60, A230, A403 and A135 and the absence of this gene was confirmed by WGS for CIP70.10, A37, A135 and shown to be truncated for A60. Studies have shown that the *adeABC* operon is not always present in *A. baumannii* strains and amongst the *adeRS-AB-*expressing strains, the outer membrane compartment gene *adeC* is not always present.^33,54^

AdeI was not detectable in any of the samples processed despite WGS confirmation. This may suggest (along with the non-detectable AdeBC mentioned above) that these proteins are not expressed in abundance in these particular isolates or a more membrane-specific sample preparation is required for better resolution of membrane proteins, although Yoon and colleagues also reported AdeB to be undetectable in parent strain BM4587 by membrane sample LC-MS/MS.^52,55^

There were no changes in abundance of AdeA, J and K in the carbapenem resistant isolates compared to CIP70.10 (**Figure 2e-g**). However, there were higher levels in one or more of the proteins in susceptible isolates A60, A230 and A90, with the former two having raised meropenem MICs of 2 mg/L. This suggests that overexpression of these efflux pumps may play a minor role in elevating MICs but the key driver of carbapenem resistance in the clinical isolates under study is the upregulation of OxaAb variants with specific amino acid substitutions.

#### (iii) Other proteins involved in membrane integrity

Changes in expression levels of penicillin-binding proteins (PBPs) has been associated with carbapenem resistance, such as the decrease in PBP2 expression levels^56^ or increase in PBP1a and 5 in an imipenem-resistant MDR strain in the presence of imipenem.^20^ Four PBPs were identified in the LC-MS/MS data – PBP1a, 2, 5 and 6 but no differences were observed between carbapenem susceptible and resistant isolates.

#### (iv) β-lactamases

Seven ADC and four OxaAb proteins encoded by genes that have upstream IS elements were detected and confirmed by WGS (**Figure 2h, 2i, Table 1**). ADC variants in CIP70.10, A90, B1 and A135 did not have an upstream IS element and this was associated with enzyme levels below the level of detection. The ADC enzyme in A230 was the only variant with IS*Aba125* upstream and displayed the lowest abundance amongst the seven variants with an IS element upstream. This implies that the promoters carried in the other IS elements are stronger than in IS*Aba125*. Only the *oxaAb* genes in carbapenem-resistant isolates A403, A371, A443 and A404 carried upstream IS elements and only these had detectable levels of enzyme. Again, this confirms that the IS elements are upregulating carbapenemase production.

### Concluding Remarks

During the course of this study, other groups published work including clinical isolate characterization and structural studies that identified residues Trp-222, Ile-129, Pro-130 and Leu-167 in the active site of OxaAb enzymes to contribute to weak carbapenem binding by obstructing the active site from carbapenem interaction, and that substitutions at these sites improve carbapenemase activity.^26,28,29^ While none of the enzymes in this study had Trp-222 or Pro-130 substitutions, this work confirms that OxaAb variants with I129L and L167V substitutions do confer raised carbapenem MICs relative to wild-type genes when all are expressed from the same promoter. When these enzymes with increased carbapenemase activity are driven by the promoter within IS*Aba1*, this confers carbapenem resistance. This was seen in recombinants lacking additional resistance proteins, and also in the resistant clinical isolates, where no additional protein abundance changes predicted to influence carbapenem MIC were observed. Hence, we conclude that overproduction of OxaAb variants with enhanced carbapenemase activity is sufficient to confer carbapenem resistance in *A. baumannii*.

## ACKNOWLEDGEMENTS

Genome sequencing was provided by MicrobesNG (http://www.microbesng.uk), which is supported by the BBSRC (grant number BB/L024209/1).

We would like to thank Laurent Poirel for kindly donating us the strains CIP70.10 and BM4547 and Te-Li Chen for providing us with vector pYMAb2.

## FUNDING

This work was funded by Anglia Ruskin University and grants MR/N013646/1 and NE/N01961X/1 from the Antimicrobial Resistance Cross Council Initiative supported by the seven United Kingdom research councils and the National Institute for Health Research.

## TRANSPARENCY DECLARATIONS

None for all authors.

## Notes

### Competing Interest Statement

The authors have declared no competing interest.

