## Supplementary material for "Variability in carbapenemase activity of intrinsic OxaAb (OXA-51-like) beta-lactamase enzymes in *Acinetobacter baumannii*": Table S1

**[1]Department of Biomedical and Forensic Science, Anglia Ruskin University, Cambridge, UK**

**[2]School of Cellular and Molecular Medicine, University of Bristol, UK**

**[3]Division of Infection & Immunity, Faculty of Medical Sciences, University College London, UK**

**[4]Bristol Proteomics Facility, University of Bristol, Bristol. UK.**

**[5]School of Biomedical Sciences, Faculty of Health, University of Plymouth, UK**

**[6]Norwich Medical School, University of East Anglia, UK**

***Correspondence: Yuiko Takebayashi, School of Cellular and Molecular Medicine, Biomedical Sciences Building, University Walk, Bristol, UK, BS8 1TD.. +44(0)117 331 2037.**

**Table S1**. **Amino acid and nucleotide changes for all OxaAb sequences at Ile-129 and Leu-167.**

**Changes at Ile-129**

| OxaAb | consensus a.a. | resulting a.a. | consensus codon | resulting codon |
|---|---|---|---|---|
| (523) | I | F | ATT | TTT |
| (110) | I | L | ATT | CTT |
| (83) | I | L | ATT | CTT |
| (312) | I | L | ATT | CTT |
| (425) | I | L | ATT | CTT |
| (516) | I | L | ATT | CTT |
| AM690768 | I | L | ATT | CTT |
| (202) | I | M | ATT | ATG |
| (248) | I | M | ATT | ATG |
| (95) | I | M | ATT | ATG |
| (131) | I | V | ATT | GTT |
| (172) | I | V | ATT | GTT |
| (173) | I | V | ATT | GTT |

**Changes at Leu-167**

| OxaAb | consensus a.a. | resulting a.a. | consensus codon | resulting codon |
|---|---|---|---|---|
| (521) | L | I | CTG | ATA |
| (127) | L | Q | CTG | CAG |
| (545) | L | V | CTG | GTA |
| NZ_LQRQ01000007 | L | V | CTG | GTG |
| NZ_KK736155 | L | V | CTG | GTG |
| (107) | L | V | CTG | GTG |
| (108) | L | V | CTG | GTG |
| (113) | L | V | CTG | GTG |
| (115) | L | V | CTG | GTG |
| (138) | L | V | CTG | GTG |
| (174) | L | V | CTG | GTG |
| (175) | L | V | CTG | GTG |
| (176) | L | V | CTG | GTG |
| (177) | L | V | CTG | GTG |
| (82) | L | V | CTG | GTG |
| (201) | L | V | CTG | GTG |
| (219) | L | V | CTG | GTG |
| (194) | L | V | CTG | GTG |
| (195) | L | V | CTG | GTG |
| (196) | L | V | CTG | GTG |
| (197) | L | V | CTG | GTG |
